# A zebrafish model of Bethlem myopathy reveals CaV1.1 as the missing link between collagen type VI deficiency and muscle dysfunction

**DOI:** 10.1101/2025.06.02.657388

**Authors:** Romane Idoux, Chloé Exbrayat-Heritier, Adrien Ducret, Shivashakthi Shivaraman, Marilyne Malbouyres, Francisco Jaque-Fernandez, Christine Berthier, Frédéric Sohm, Vincent Jacquemond, Sandrine Bretaud, Florence Ruggiero, Bruno Allard

**Author notes:** Co-corresponding authors: Florence Ruggiero and Bruno Allard. These authors contributed equally to this work. **Email:** and. **Author Contributions:** Conceptualization: RI, SB, FR, BA; Methodology: RI, SB, CB, AD, FR, BA; Investigation: RI, CE-H, FJ-F, SS, CB; Supervision: SB, FS, FR, BA; Writing—original draft: FR, BA; Writing—review & editing: VJ, FR, BA. **Competing Interest Statement:** The authors do not declare any competing interest.

## Abstract

Bethlem myopathy (BM) results from mutations in genes encoding one of the three α chains of collagen VI (ColVI). This muscle disease is characterized by skeletal muscle weakness and wasting worsening with age. How alteration in ColVI present outside muscle fibers in the extracellular matrix induces dysfunction within muscle fibers is still misunderstood. Here we explored intracellular Ca^2+^ handling properties in isolated fast skeletal muscle fibers from adult zebrafish harboring a mutation (*col6a1*^*Δex14*^) that is the most frequently found in BM patients. *Col6a1*^*Δex14*^ fish muscle exhibited progressive loss of ColVI deposition, defects in basement membrane and ColVI intracellular accumulation. By combining voltage-clamp and intracellular Ca^2+^ measurements on isolated fibers, we showed that voltage-dependence of intramembrane charge movements produced by the activation of CaV1.1 controlling sarcoplasmic reticulum (SR) Ca^2+^ release and voltage-dependence of depolarization-induced SR Ca^2+^ release were shifted toward negative potentials in *col6a1*^*Δex14*^ fish. These changes in voltage-dependence gave rise to larger SR Ca^2+^ leak at voltages close to resting values and to higher frequency of spontaneous SR Ca^2+^ release elementary events in mutant fish. Trunk muscle force and swimming performance were also found to be reduced in *col6a1*^*Δex14*^ fish and mis-localization of CaV1.1 subunits clusters was observed in mutant fibers t-tubules. These data indicate that ColVI deficiency in BM leads to CaV1.1 dysfunction that contributes to promote a pathogenic SR Ca^2+^ leak responsible for progressive muscle weakness and wasting. CaV1.1 could represent the still elusive transmembrane link allowing altered myomatrix to transduce pathogenic signals within muscle.

## Introduction

Bethlem myopathy (BM) is a debilitating genetic disorder characterized by abnormal contractures of the Achilles tendons, elbows, shoulders, spine and finger flexors, experienced as joint stiffness and associated with proximal muscle weakness. Progressive muscle wasting also occurs and symptoms worsen in the 4^th^ and 5^th^ decades of life with increasing risk of respiratory insufficiency and ambulation difficulties (1-3). The pathology is incurable and its prevalence is 0.77 per 100,000. BM is an autosomal dominant disease caused by mutations in the *COL6A1, COL6A2, COL6A3* genes that encode the three major chains of collagen type VI (ColVI)-, α1, α2, α3 respectively. ColVI is a component of the extracellular matrix (ECM) with a broad tissue distribution including high expression in skeletal muscle. In skeletal muscle, ColVI is produced by interstitial fibroblasts where three ColVI α-chains assemble into triple-helix monomers, themselves assembling into dimers, then tetramers that are secreted in the extracellular space to associate into characteristic beaded microfilaments (4, 5). ColVI binds to a large number of ECM components, structuring in this way the three-dimensional architecture of the tissue and connecting directly or indirectly muscle cells to ECM (5). Nowadays, a still unresolved issue is to understand how a disrupted ColVI protein belonging to ECM can induce pathologic muscle dysfunction?

In an attempt to address this question, some animal models were generated such as a mouse and a zebrafish model in which the *col6a1* or *col6a2* gene has been invalidated (6-8). Expression of different ColVI α chains has been also transitory knocked down by morpholinos injection in zebrafish larvae (9). Among these models, several muscle abnormalities have been reported such as a loss of contractile strength, sarcoplasmic reticulum (SR) alterations, neuromuscular transmission deficiency and structural and functional impairment of mitochondria and dysregulation of the TGF-β pathway (7-11). However, microinjection of morpholinos in developing embryos provokes transient knockdown of targeted genes while BM is a progressive disease. Moreover, invalidation of *col6a1* in mice provokes a complete loss of ColVI production, whereas in humans with BM mutations in any of the three genes rarely cause ColVI complete loss.

In order to circumvent these limitations, Radev *et al*.(12) created the first zebrafish line harboring an in-frame exon deletion that is the most frequent mutation found in BM patients. Using a transcription activator-like effector nuclease (TALEN) strategy, they provoked a deletion in a splice donor site leading to an in-frame skipping of exon 14 in *col6a1* mRNA (*col6a1*^*Δex14*^). As described in *Col6a1*^*-/-*^ mouse and morpholino-knock down zebrafish models, ultrastructural analysis of zebrafish *col6a1*^*Δex14-/-*^ (hereafter referred to *col6a1*^*Δex14*^) trunks at 6-7 dpf, 1 week and 1 year revealed the same ultrastructural abnormalities, including SR alteration, which worsen with age (12) as observed in the human disease.

Interestingly, it was found that intracellular Ca^2+^ increase provoked by inhibition of the mitochondrial F_1_-F_0_ATPase was larger and occurred earlier in muscle from *Col6a1*^*-/-*^ as compared to wild-type (WT) mice and was suppressed by dantrolene, a blocker of SR Ca^2+^ release channel (10), suggesting a possible dysfunction of SR Ca^2+^ flux associated with SR structure alterations as observed in mice and in *col6a1*^*Δex14*^ zebrafish. In skeletal muscle, Ca^2+^ ions activating contraction are released by the SR in response to muscle excitation through the excitation-contraction (EC) coupling process. During this process, action potentials activate the skeletal muscle voltage-gated Ca^2+^ channel (CaV1.1), resulting in intramembrane charge movements, which in turn opens the ryanodine receptor (RyR_1_), a Ca^2+^ release channel anchored in the SR membrane (13). We recently provided a comprehensive characterization of EC coupling properties of isolated fast muscle fibers from adult zebrafish and demonstrated that zebrafish EC coupling displays superfast properties (14). The reported effect of dantrolene and altered SR structure in *col6a1* deficient muscle fibers, together with the fact that BM symptoms worsen with age, prompted us to explore in the present study the properties of EC coupling using current- and voltage-clamp combined with intracellular Ca^2+^ measurement in isolated fibers from *col6a1*^*Δex14*^ zebrafish at adult stage. Additionally, we characterized the defects in synthesis, secretion, oligomerization and expression of ColVI α1 chains in muscles from adult *col6a1*^*Δex14*^ zebrafish. Our data give evidence of an alteration of the functional expression and of the voltage-dependence of CaV1.1 that contributes to promote a pathogenic SR Ca^2+^ leak in *col6a1*^*Δex14*^ zebrafish, and to severely affect muscle physiology, as evidenced by the observed defects in muscle performance and swim behavior.

## Results

### Defects in basement membrane and ColVI intracellular accumulation in mutant muscle

The exon 14 deletion in *col6a1* predicts a truncated ColVI α1 chain (12), which may interfere with trimer formation alongside its two partner chains, α2(VI) and α3(VI). To assess trimer assembly and secretion of ColVI, we performed immunofluorescence staining using guinea pig antibodies raised against the recombinant C-terminal domain of the zebrafish α1(VI) chain (see Methods) on frozen muscle sections of 6-month-old and 1-year-old mutant and wild-type (WT) zebrafish (Fig. 1). In WT fish, a continuous ColVI staining outlining all muscle fibers is observed (Fig. 1A, left panel). In striking contrast, *col6a1*^*Δex14*^ mutants show an interrupted, patchy ColVI signal surrounding muscle cells, with areas where ColVI is absent (Fig. 1A, right panel). This suggests that the production of the truncated α1(VI) chain by muscle fibroblasts impairs the extracellular ColVI filament assembly. ColVI immunoreactivity is nearly absent in 1-year-old wild-type muscle, mirroring the age-related decline in BM patients (Fig. 1A). However, ColVI staining varied, showing either a complete absence or the presence of a few residual patches depending on the individual (Fig. S1). ColVI is a basement membrane-associated collagen (5). We therefore examined whether disorganized ColVI or its age-dependent loss in 1-year-old muscle affects laminin deposition, a major basement membrane component. Immunofluorescence with pan-laminin antibodies revealed that laminin deposition is disrupted in mutant muscle, presenting as a patchy pattern at both 6 months and 1 year compared to the continuous fluorescent line delineating each myofiber in WT muscle (Fig. 1A). Unlike ColVI, laminin remained detectable with age, although its organization was clearly disturbed. ColVI immunoreactivity was further confirmed on isolated muscle cells from 1-year old zebrafish trunks (Fig. 1B). This allowed the observation of an intracellular retention of ColVI in neighboring fibroblasts, the primary source of ColVI in skeletal muscle, indicating that although these cells still produce the mutant protein, its secretion is impaired. Western blot analysis on protein extracts from trunk muscles confirmed the immunofluorescence data showing a faint α1(VI) chain band in mutant samples, in contrast to WT muscles (Fig. 1C and densitometry). In conclusion, ColVI deposition is disrupted in the ***col6a1***^***Δex14***^ mutant fish, characterized by at best a patchy presence of the protein around skeletal muscle fibers. This disruption affects basement membrane assembly and may impair its mechanical and/or signaling roles in mutant skeletal muscle.

**Fig. 1.**
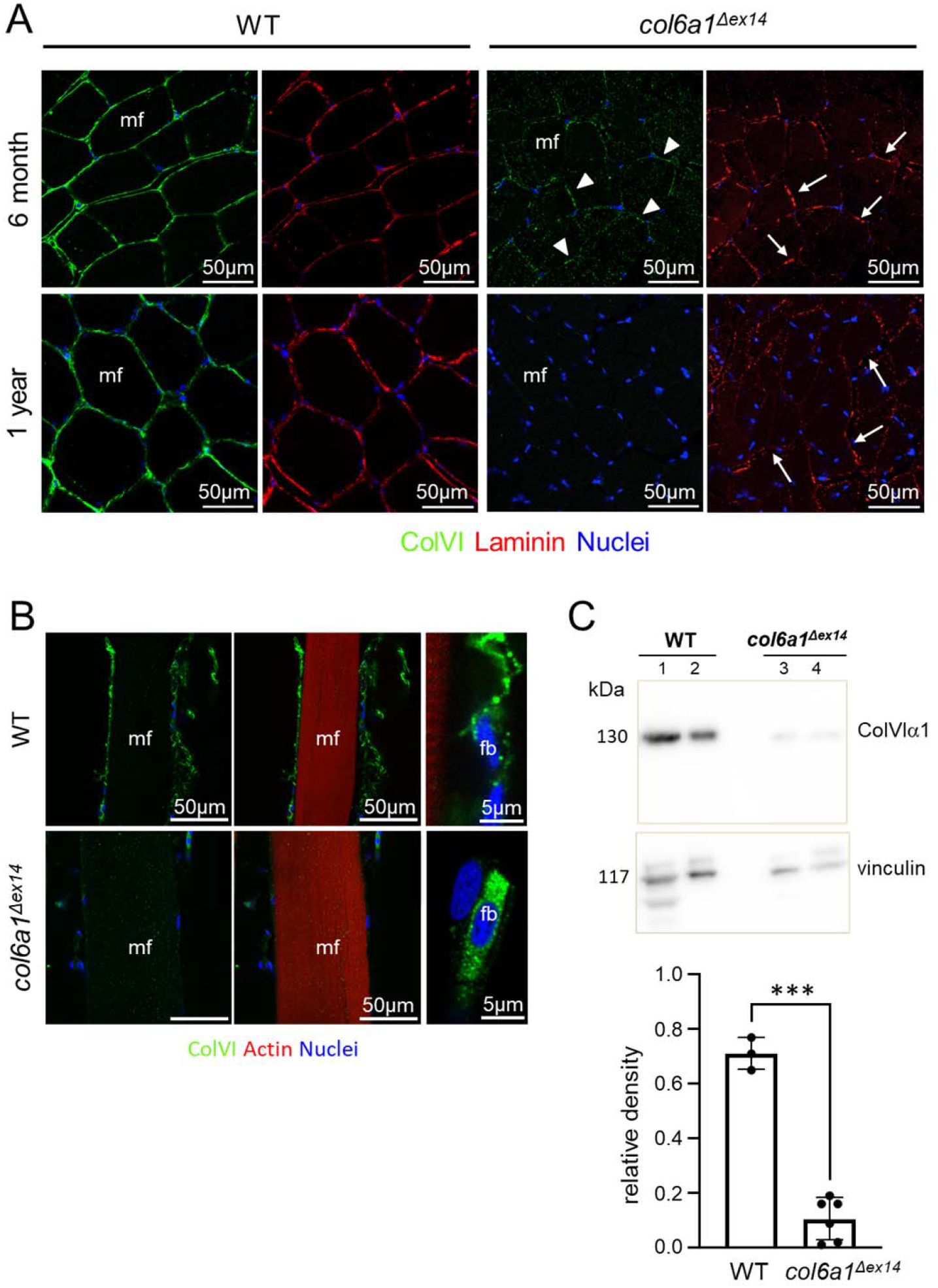
ColVI deposition, basement membrane organization and ColVI intracellular accumulation in *col6a1*^*Δex14*^ mutant. (A) Confocal images of 6-month and 1-year old zebrafish muscle cross-section (mf, muscle fibers; arrowheads, aggregates; arrow, interruption. (B) Confocal images of 1-year old zebrafish isolated muscle fibers (mf, muscle fiber; fb, fibroblast). (C) Western blotting with anti-ColVI antibodies of 1-year old muscle extracts. Vinculin antibodies were used as loading controls. Data from one representative experiment are shown. Histogram shows densitometric quantification of relative ColVIa1 expression levels normalized to vinculin. Band intensities were quantified using ImageJ software and a paired t-test was used to determine statistical significance. Data were obtained in 3 WT and 6 *col6a1*^*Δex14*^ fish.

### T-tubule structure is not altered in mutant muscle fibers

In skeletal muscle, excitation-contraction coupling involves two membrane systems, the transverse tubules (t-tubules) and the SR. The t-tubules correspond to extensions of the sarcolemma that penetrate into the depth of the fiber, allowing action potentials propagation. In order to investigate potential alterations in the structure of t-tubules, isolated WT and *col6a1*^*Δex14*^ muscle fibers were stained with di-8-anepps. Fig 2A shows confocal images of di-8-anepps fluorescence in a WT fiber (left) and a *col6a1*^*Δex14*^ fiber (right). In both fiber populations, the fluorescence patterns were comparable, displaying transverse striations consistent with T-tubule localization at Z-lines of sarcomeres in zebrafish muscle. No T-tubule misorientation was detected in the *col6a1*^*Δex14*^ group, and the T-tubule organization index did not differ significantly between WT and *col6a1*^*Δex14*^ (mean T-tubule power ± SD was 89.9 ± 6.5 in WT *versus* 85.98 ± 9.45 in mutant) (Fig. 2B). Likewise, longitudinal T-tubule spacing, corresponding to the interval between t-tubules and reflecting sarcomere length, was similar in the two groups (mean period ± SD was 1.83 ± 0.02 μm in WT and 1.84 ± 0.03 μm in mutant) (Fig. 2C). These data indicate that T-tubule organization is preserved in *col6a1*^*Δex14*^ zebrafish.

**Fig. 2.**
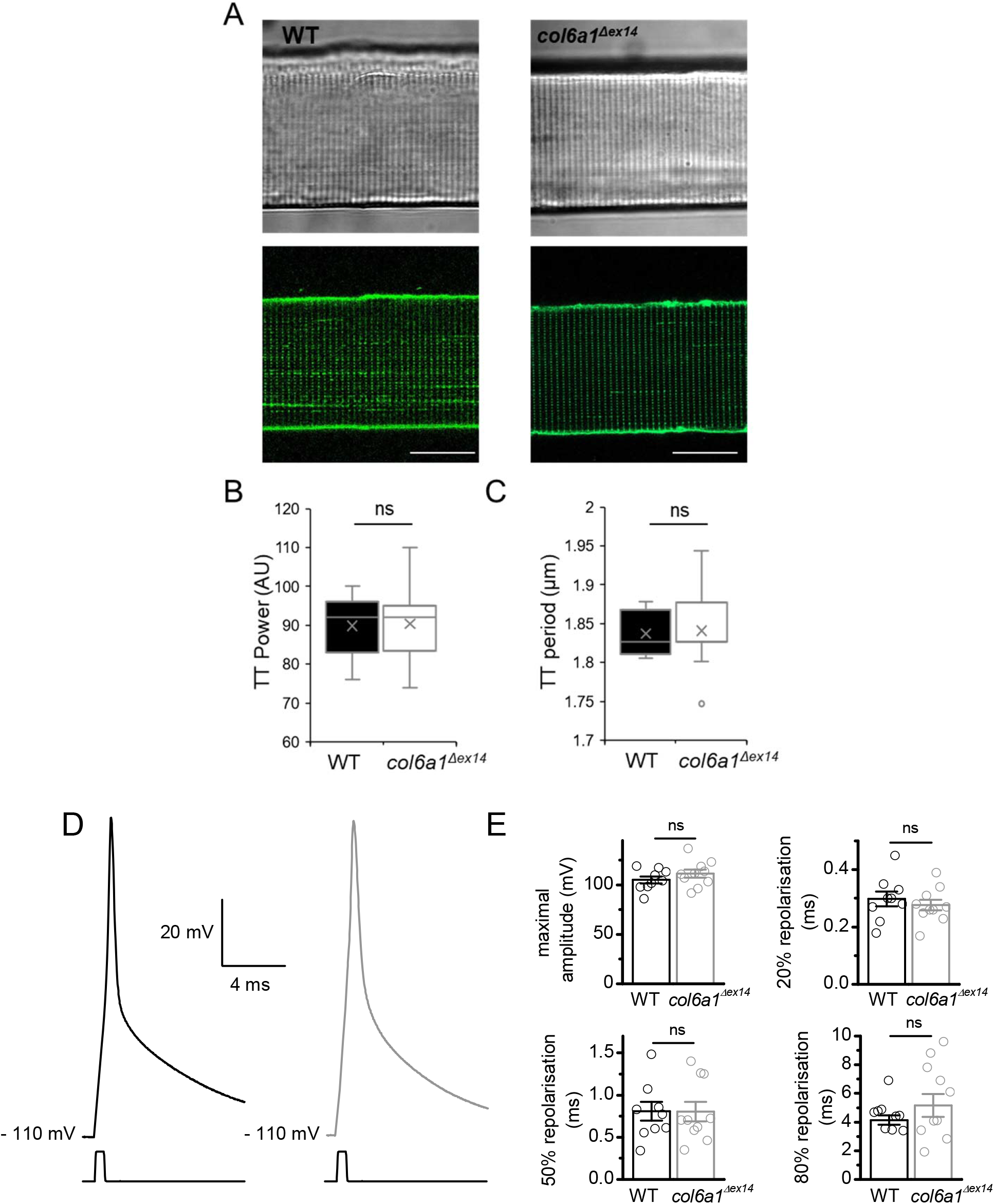
T-tubules network and action potentials in WT and *col6a1*^*Δex14*^ muscle fibers. (A) Representative transmitted light and x, y fluorescence confocal images of a WT (left) and a *col6a1*^*Δex14*^ (right) isolated muscle fiber stained with di-8-anepps. Scale bar = 20 μm. (B) Longitudinal spacing regularity of the T-tubular network (T-tubule power) in WT and *col6a1*^*Δex14*^ muscle fibers. (C) Mean longitudinal t-tubules spacing (TT period) in WT and col6a1^Δex14^ muscle fibers. Image analysis was performed on two confocal sections per fiber, yielding a total of 44 images from 22 fibers of three 1-year-old WT fish, and 46 images from 23 fibers of three 1-year-old *col6a1*^*Δex14*^ fish. Box plots show the interquartile range (box, 50% of the values), the median (line), the mean (cross), and outliers (circles). No statistically significant differences were detected between WT and *col6a1*^*Δex14*^ *fibers*, as assessed by the Mann– Whitney–Wilcoxon test. (D) Action potential elicited by injection of a 0.5-ms-long suprathreshold depolarizing current in current clamp condition in a WT (left) and a *col6a1*^*Δex14*^ (right) isolated muscle fibers. (E) Histograms presenting the maximal amplitude, time to peak and time for 20, 50 and 80 % repolarization of action potential after spike in WT and *col6a1*^*Δex14*^ fibers. Data were obtained in 9 fibers from 3 WT fish and 10 fibers from 2 *col6a1*^*Δex14*^ fish.

### Action potentials properties are not affected in mutant muscle fibers

Under physiological conditions, the first step of EC coupling corresponds to the firing and propagation of action potential along sarcolemma and t-tubules. A series of current-clamp experiments were performed to measure action potentials in WT and *col6a1*^*Δex14*^ fibers. In both fiber populations, injection of 0.5-ms duration suprathreshold depolarizing current from a resting membrane potential held at -110 mV and in the presence of an external Tyrode solution induced action potentials (Fig. 2D). In WT muscle, as described previously, the peak value of voltage reached by action potentials was close to 0 mV and the repolarization phase displayed very fast kinetics (14). Fig 2E shows that maximal amplitude, time to peak and time for 20, 50 and 80 % repolarization of action potential after spike were not significantly different in WT and *col6a1*^*Δex14*^ muscle fibers. These data suggest that membrane excitable properties are not affected in *col6a1*^*Δex14*^ fish muscle.

### Alteration of density and of voltage dependence of charge movements in mutant muscle fibers

A next series of experiments aimed at studying the functionality of CaV1.1 that controls in a voltage-dependent manner the opening of RyR_1_ channels, the SR Ca^2+^ channels responsible for the intracellular Ca^2+^ release. For this purpose, the membrane currents resulting from the movement of intramembrane charges carried by CaV1.1 were recorded in response to depolarization. Figure 3A shows currents produced by intramembrane charge movements recorded in a voltage-clamped zebrafish muscle fiber in the absence of external Na^+^ and K^+^ ions and in the presence of Na^+^ and K^+^ channel blockers and low Cl^-^ from a holding potential of -80 mV in WT and *col6α1*^*Δex14*^ muscle fiber. The density of “on” charges (Q_on_) was plotted as a function of voltage and the relationship was fitted in each cell with a Boltzmann equation (Fig. 3B). The mean values for Q_max_, V_0.5_, and k were 8.8 ± 0.7 nC/μF, −32.5 ± 1.5 mV, and 9 ± 0.4 mV for WT fish and 6.3 ± 0.7 nC/µF, -41.4 ± 1.4 mV, and 6.7 ± 0.4 mV for *col6a1*^*Δex14*^ fish respectively (Fig. 3C). The maximal charges density (Q_max_) was significantly reduced in *col6a1*^*Δex14*^ fibers indicating a lower density of functional CaV1.1 in *col6a1*^*Δex14*^ fish (P= 0.027). The voltage dependence of charge movements was also significantly steeper and shifted toward negative potentials in *col6a1*^*Δex14*^ fish.

**Figure 3.**
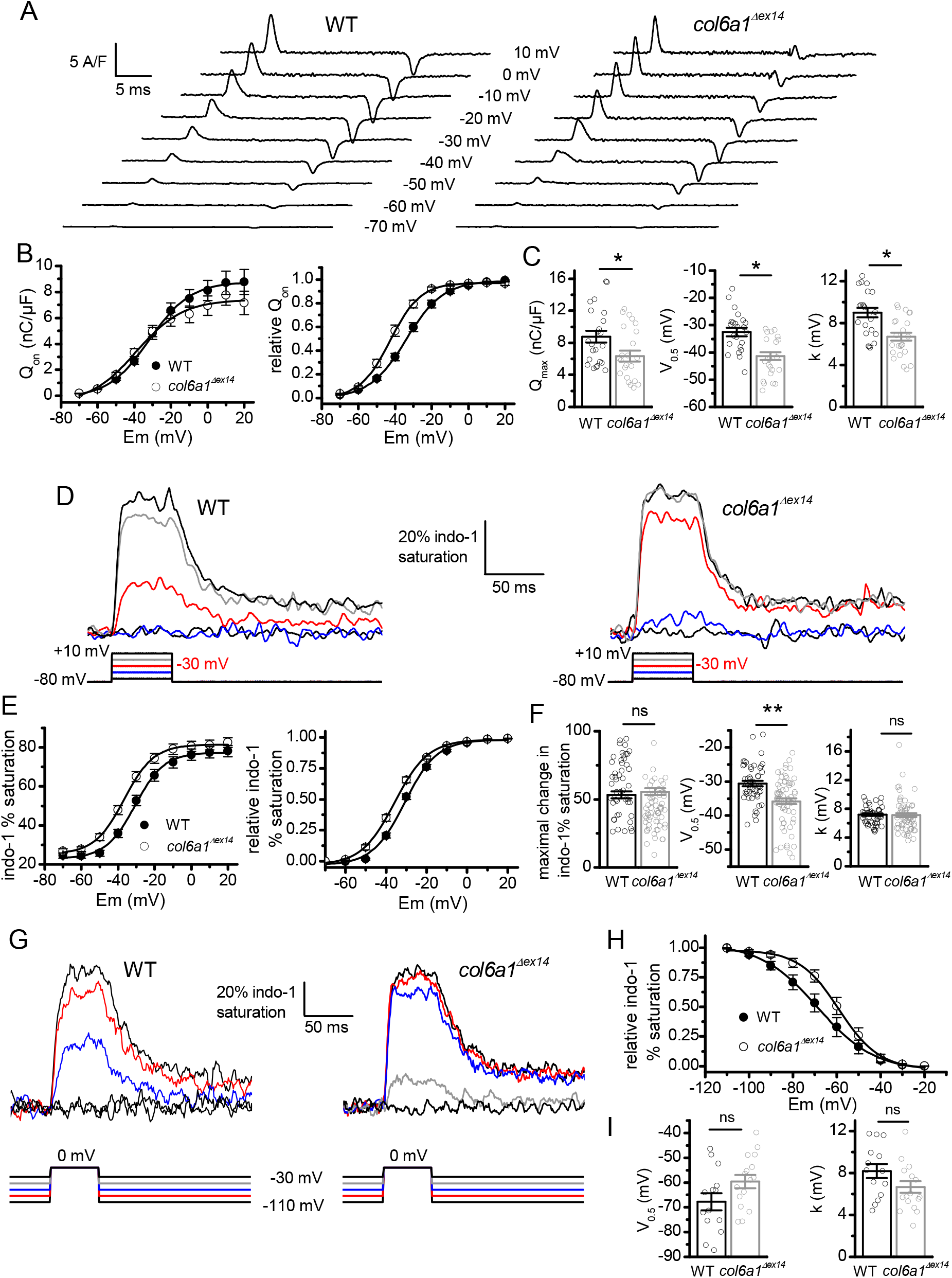
Intramembrane charge movements and voltage-activated Ca^2+^ transients in WT and *col6a1*^*Δex14*^ muscle fibers. (A) Charge movements recorded in response to 20-ms-long depolarizing pulses from a holding potential of -80 mV to the indicated value in a WT (left) and in a *col6a1*^*Δex14*^ (right) isolated muscle fiber. (B) Relationship between the mean density of charge movements and membrane potential in WT and *col6a1*^*Δex14*^ fibers before (left) and after normalization of charge movements density (right). In the left graph, mean data points from the 2 curves were fitted using a Boltzmann equation with values for Q_max_, V_0.5_ and k of 8.8 nC/μF, -32 mV and 11 mV in WT fibers respectively, and of 7.4 nC/μF, -39 mV and 13 mV in *col6a1*^*Δex14*^ fibers respectively. In the right graph, mean data points from the 2 curves were fitted using a Boltzmann equation with values for V_0.5_ and k of -34 mV and 10 mV in WT fibers respectively, and of -43 mV and 8 mV in *col6a1*^*Δex14*^ fibers respectively. (C) Histograms presenting mean values of Q_max_, V_0.5_ and k in WT and in *col6a1*^*Δex14*^ fibers. Data were obtained in 23 fibers from four WT fish and 24 fibers from three *col6a1*^*Δex14*^ fish. (D) Indo-1 percentage saturation traces in response to 50-ms-long depolarizing pulses of increasing amplitude in a WT (left) and a *col6a1*^*Δex14*^ (right) isolated muscle fiber. Traces have been smoothed using adjacent averaging of 20 datapoints. (E) Relationship between the maximal amplitude of indo-1 percentage saturation and the membrane potential in WT and *col6a1*^*Δex14*^ fibers before (left) and after normalization of indo-1 percentage saturation (right). In the left graph, mean datapoints from the two curves were fitted using a Boltzmann equation with values for maximal indo-1 % saturation, V_0.5_ and k of 77, -31 mV and 8 mV in WT fibers respectively, and of 81, -37 mV and 8 mV in *col6a1*^*Δex14*^ fibers respectively. In the right graph, mean data points from the 2 curves were fitted using a Boltzmann equation with values for V_0.5_ and k of - 31 mV and 8 mV in WT fibers respectively, and of -36 mV and 8 mV in *col6a1*^*Δex14*^ fibers respectively. (F) Histograms presenting mean values of maximal change in indo-1 % saturation from resting value, V_0.5_ and k in WT and *col6a1*^*Δex14*^ fibers. Data were collected in 50 fibers from seven WT fish and 66 fibers from eight *col6a1*^*Δex14*^ fish. (G) Indo-1 signals elicited by 50-ms-depolarizing pulses at 0 mV from holding potentials (HP) increasingly depolarized to the indicated voltages in a WT (left) and a *col6a1*^*Δex14*^ (right) isolated muscle fiber. Traces have been smoothed using adjacent averaging of five datapoints. (H) Relationship between mean relative peak of indo-1 percentage saturation in response to depolarizing pulses at 0 mV and holding potentials in WT and *col6a1*^*Δex14*^ fibers. Mean data points from the two curves were fitted using a Boltzmann equation with values for V_0.5_ and k of -69 mV and 13 mV in WT fibers respectively, and of -59 mV and 9 mV in *col6a1*^*Δex14*^ fibers respectively. (I) Histograms presenting V_0.5_ (mV) and k (mV) in WT and *col6a1*^*Δex14*^ fibers. Data have been obtained in 14 fibers from two WT fish and 16 fibers from two *col6a1*^*Δex14*^ fish.

### Negative shift of voltage dependence of SR Ca^2+^ release in mutant muscle fibers

By activating CaV1.1 proteins, depolarization induces the opening of SR Ca^2+^ release channels. Using the same external solution as the one used for recording charge movements, intracellular Ca^2+^ transients were measured in response to 50-ms duration depolarizing pulses of increasing amplitudes with the ratiometric Ca^2+^ indicator indo-1 dialyzed into WT and *col6a1*^*Δex14*^ cells (Fig 3D). The indo-1 % saturation was plotted as a function of voltage and the relationship was fitted in each cell with a Boltzmann equation (Fig. 3E). Maximal change in indo-1 % saturation from resting value, V_0.5_, and k were 53.2 ± 2.6 %, -30.6 ± 0.8 mV and 7.2 ± 0.2 mV for WT fish and 55.6 ± 2.4 %, -35.8 ± 0.9 mV and 7.2 ± 0.3 mV for *col6a1*^*Δex14*^ fish respectively. Maximal amplitude of voltage-induced Ca^2+^ transients and steepness of voltage-dependence were not significantly changed in *col6a1*^*Δex14*^ fibers but the voltage dependence was significantly shifted toward negative potentials (Fig. 3F). The mean indo-1 % saturation in fibers held at -80 mV was found to be not significantly changed in *col6a1*^*Δex14*^ fibers (26.8 ± 0.01 %) as compared to WT fibers (25.8 ± 0.01 %).

We showed in a previous study that prolongation of depolarization induces a progressive decline of the amplitude of Ca^2+^ transients resulting from a time- and voltage-dependent inactivation of CaV1.1 (14). This inactivation process was investigated in WT and *col6a1*^*Δex14*^ by applying 50-ms duration voltage pulses to 0 mV from holding potentials incrementally depolarized during 20 s (Fig. 3G). The relative amplitude of the Ca^2+^ transients was plotted as a function of the holding potential in each cell and the relationships were fitted with a Boltzmann equation (Fig. 3H). Values for V_0.5_ and k were -67.7 ± 3.4 mV and 8.2 ± 0.7 mV for WT fish and -59.6 ± 2.7 mV and 6.7 ± 0.6 mV for *col6a1*^*Δex14*^ fish respectively (Fig. 3I). Although the relationship between Ca^2+^ release inactivation and voltage in *col6a1*^*Δex14*^ fish tended to be shifted toward positive potentials, none of these parameters were significantly different.

### Low amplitude voltage pulses induce larger intracellular Ca^2+^ changes in mutant muscle fibers

The negative shift of the voltage activation of charge movements and SR Ca^2+^ release observed in *col6a1*^*Δex14*^ fiber suggests that opening of RyR_1_ may occur at membrane potentials more negative than the voltage threshold of Ca^2+^ release usually observed between -50 and -40 mV in WT muscle (13, 14). The use of 50-ms duration depolarizing pulses together with global fluorescence measurements in the preceding experiments did not provide the required resolution for exploring intracellular Ca^2+^ changes provoked by low values of depolarization. Hence, in order to assess possible changes in voltage sensitivity of CaV1.1 at membrane potentials more negative than -50 mV, we measured intracellular Ca^2+^ changes in response to long duration (10-s) depolarizing pulses from a holding potential of -90 mV using Fluo-4 fluorescence confocal imaging in WT and *col6a1*^*Δex14*^ fibers. Figure 4A shows that Ca^2+^ changes induced by depolarizing pulses from -80 to - 60 mV were significantly larger in the *col6a1*^*Δex14*^ fiber. In average, plotting the changes in fluorescence as a function of depolarization indicated that the maximal Ca^2+^ level reached during the pulses was significantly higher for all depolarization pulses from -80 to -60 mV in *col6a1*^*Δex14*^ (Fig. 4B). In addition, it can be observed in Fig. 4A that Ca^2+^ remained at a larger level during the 40-s interval of repolarization between pulses in the *col6a1*^*Δex14*^ fiber. In average, plotting the changes in the basal fluorescence before each pulse as a function of depolarization showed that the basal Ca^2+^ increased at a significantly higher level with consecutive pulses in *col6a1*^*Δex14*^ fibers (Fig. 4B).

**Figure 4.**
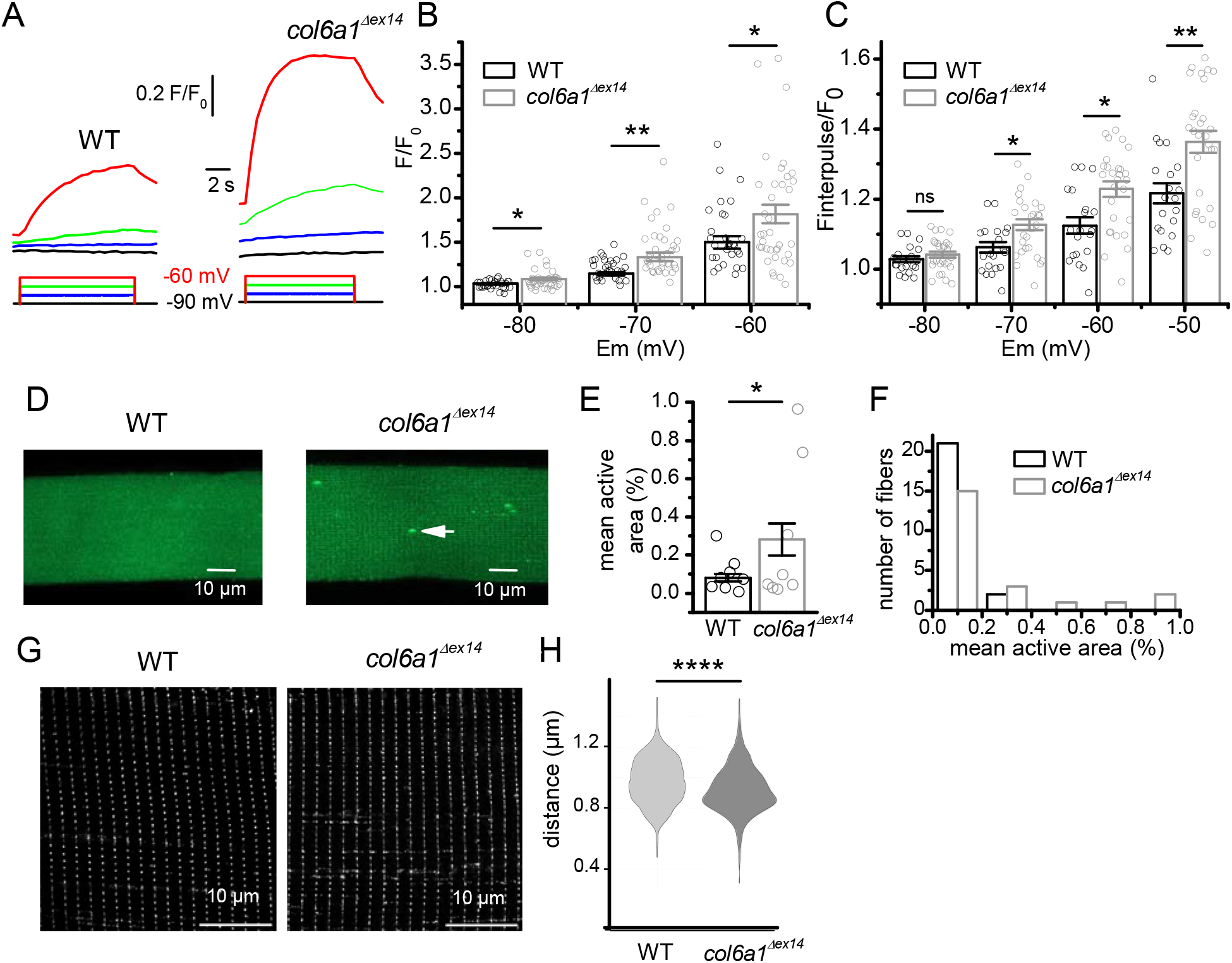
Intracellular Ca^2+^ changes evoked by long duration depolarizing pulses of low amplitude, Ca^2+^ sparks activity, and DHPR distribution along t-tubules in WT and *col6a1*^*Δex14*^ muscle fibers. (A) Fluo-4 fluorescence changes expressed as F/F_0_, recorded at a frequency of 2 Hz in response to 10-s-long depolarizing pulses of increasing amplitude delivered every 50 s from a holding potential of -90 mV in a WT (left) and a *col6a1*^*Δex14*^ (right) isolated muscle fiber, where F and F_0_ correspond to the maximum fluorescence measured during the pulse and F_0_ to the fluorescence measured at -90 mV at the beginning of the experiment, respectively. (B) Mean maximal amplitude of F/F_0_ as function of membrane potential in WT and *col6a1*^*Δex14*^ fibers. Data have been obtained in 29 fibers from five WT fish and in 37 fibers from five *col6a1*^*Δex14*^ fish. (C) Mean F_interpulse_/F_0_ as function of pulse voltage in WT and *col6a1*^*Δex14*^ fibers, where F_interpulse_ corresponds to the fluorescence measured at -90 mV before the pulse. Data have been obtained in 20 fibers from four WT fish and in 28 fibers from four *col6a1*^*Δex14*^ fish. (D) Confocal fluo-4 fluorescence images during a sequence of 40 successive frames in a WT fiber and in a *col6a1*^*Δex14*^ isolated muscle fiber. Arrow points to a Ca^2+^ sparks. (E) Histogram presenting mean active area (displaying Ca^2+^ sparks) in WT and *col6a1*^*Δex14*^ groups. (F) Histograms presenting the number of WT and *col6a1*^*Δex14*^ fibers displaying increasing range of active area. Data have been obtained in 23 fibers from two WT fish and in 23 fibers from two *col6a1*^*Δex14*^ fish. (G) Airyscan detector and processing of fluorescence confocal images of 1-year old zebrafish isolated muscle fibers stained with antibodies to DHPR α1s subunit. (H) Distances between immunofluorescent foci (900 length measurements in four WT and four mutant fish). Statistics were performed using the Brunner–Munzel test, an adaptation of the Mann–Whitney U test that does not assume equal variances or a pure location shift.

### Higher Ca^2+^ sparks occurrence in resting conditions in mutant muscle fibers

The shifted activation of SR Ca^2+^ release toward negative voltages might suggest that the active control of CaV1.1 on gating of RyR_1_channel demonstrated in resting skeletal muscle (15) is impaired in *col6a1*^*Δex14*^ fibers, leading to elevated SR Ca^2+^ efflux at rest. Elevated SR Ca^2+^ efflux induced by a loss of CaV1.1 repressive action on RyR_1_channel is known to be correlated with an increase in the occurrence of spontaneous elementary events of SR Ca^2+^ release, named Ca^2+^ sparks (16). We thus looked to such events in WT and *col6a1*^*Δex14*^ fibers loaded with Fluo4. Figure 4C shows fluorescence images of a WT and a *col6a1*^*Δex14*^ fiber in which Ca^2+^ ions were homogeneously distributed in the cytosol. No Ca^2+^ sparks were detected in the WT fiber while three Ca^2+^ sparks were detected at the same time in the *col6a1*^*Δex14*^ fiber. Quantification of the percentage of fiber area exhibiting Ca^2+^ sparks in each fiber indicated that, in average, this relative fiber area was significantly larger in *col6a1*^*Δex14*^ fibers (Fig. 4D). However, because a strong variability of Ca^2+^ sparks occurrence was observed from cell to cell, we classified WT and *col6a1*^*Δex14*^ fibers according to different ranges of active area (Fig. 4D). Comparable numbers of WT and *col6a1*^*Δex14*^ fibers were found to display any or very low active area but 4 *col6a1*^*Δex14*^ fibers and none of WT fibers displayed Ca^2+^ sparks with a relative active area between 0.4 and 1%.

### Alteration of CaV1.1 distribution along t-tubules in mutant muscle fibers

The observed changes in density of charge movements, SR Ca^2+^ release and Ca^2+^ sparks in mutant fish point out to a dysfunction of CaV1.1 resulting from ColVI deficiency. These data prompted us to explore the expression pattern of CaV1.1 proteins using immunofluorescence labelling with antibodies raised against CaV1.1 in WT and in *col6a1*^*Δex14*^ fibers. Fluorescent confocal images revealed in both fish an intracellular striated expression pattern consistent with location of CaV1.1 in t-tubules (Fig. 4E). Spacing between fluorescent striations were found to be not significantly different in WT and in *col6a1*^*Δex14*^ fibers, confirming that t-tubule periodicity and sarcomere length were not altered in mutant fish as shown in Fig. 2. As already described in myotubes from zebrafish larvae (17), transverse striations consisted in alignments of fluorescent foci corresponding to clusters of CaV1.1 at the level of t-tubule/SR junctions. The mean distance between foci was found to be significantly shorter in mutant fish (Fig. 4F) indicating possible altered distribution of CaV1.1 along t-tubules.

### Twitch muscle force and swimming performance are defective in mutant zebrafish

A last series of experiments were performed in order to determine whether the observed changes in EC coupling in *col6a1*^*Δex14*^ muscle were associated with a change in muscle performance under physiological conditions. To address this question, we first measured force developed by trunk muscle from 2 weeks post-fertilization (wpf) WT and mutant fish in response to electrical field stimulation. Figure 5A and B show that the contractile force generated by mutant muscles was significantly lower than WT and the mean amplitude of the twitch response was about one third smaller than WT. Swimming performance was also measured in 1-year old WT and *col6a1*^*Δex14*^ zebrafish. Fish were placed in a swim tunnel and challenged by increasing water flows against which fish managed to swim (Fig. 5B, left). Image captures of videos at 5 BL.s^-1^ at the indicated frames showed that mutant fish had difficulty maintaining their position in the swim tunnel compared to WT fish (Fig. 5B, left). The maximal velocity that fish could reach during the swimming step protocol expressed as U_crit_ was significantly lower in *col6a1*^*Δex14*^ as compared to WT zebrafish (6.99 ± 0.37 BL.s^-1^ for WT *vs* 8.87 ± 0.52 BL.s^-1^ for *col6a1*^*Δex14*^) (Fig. 5B, right panel).

**Fig. 5.**
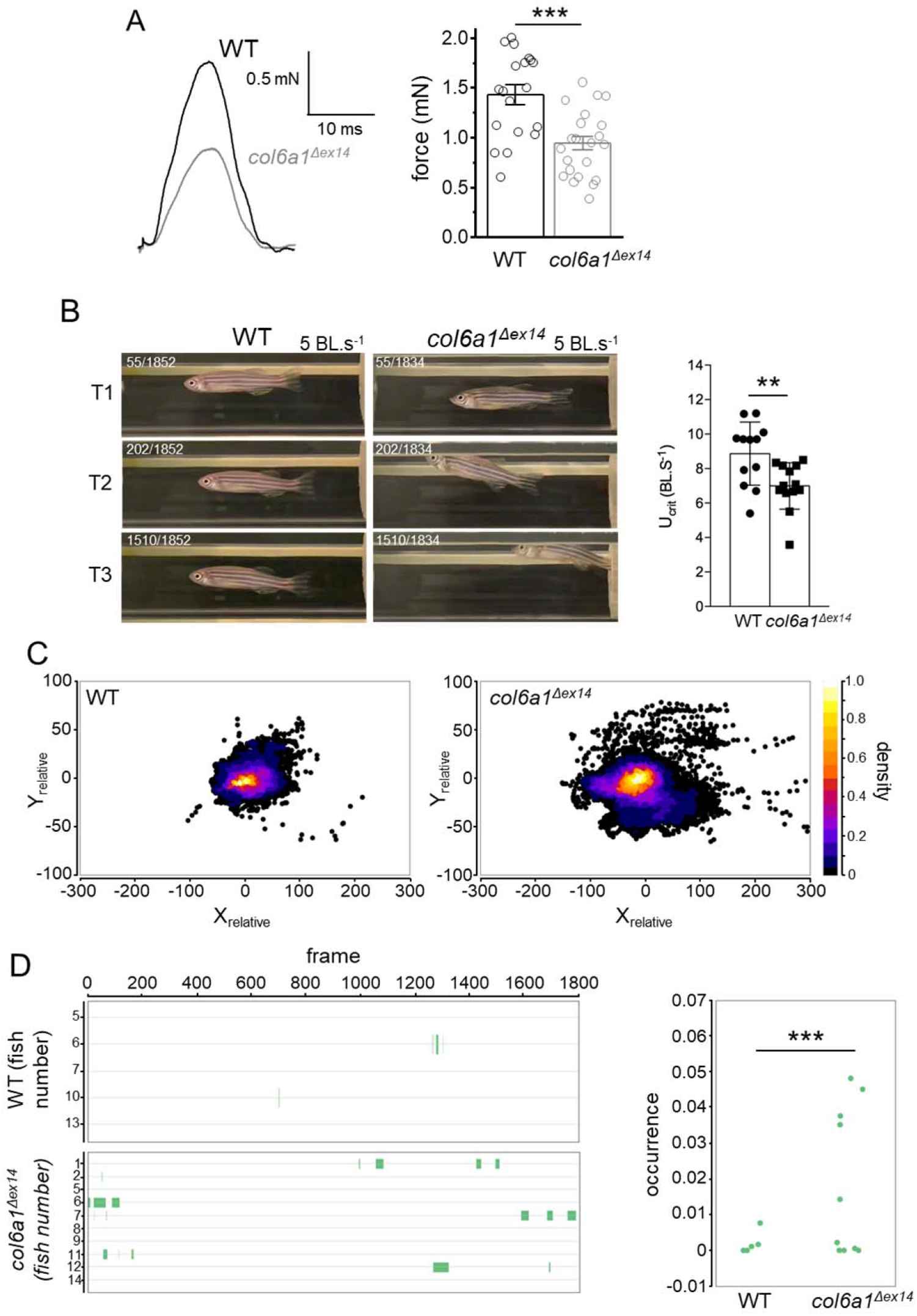
Muscle contractile force and swimming performance of WT and *col6a1*^*Δex14*^ fish. (A) Representative twitch responses elicited by single supramaximal electric shocks of 0.5 ms duration (left panel) and histograms showing mean twitch amplitudes in 19 WT and 22 *col6a1*^*Δex14*^ fish (right panel). (B) Frames from 1-min video of swimming tunnel at T1, T2 and T3, corresponding to start, middle and end, of a WT and a *col6a1*^*Δex14*^ fish at 5 BL.s^-1^ (left panel). Mean critical swimming speed (Ucrit) in 12 WT and 13 mutant fish (right panel). (C) Relative displacement of the fish. The center of the representation is the barycenter of 1-min trajectory for each WT and *col6a1*^*Δex14*^ fish (6 WT and 10 *col6a1*^*Δex14*^ fish). (D) Analysis of defective swim behavior occurrence along the X- and Y-axes in 6 WT and 10 *col6a1*^*Δex14*^ fish (left panel) and plot of occurrence values (right panel) (U-statistic).

In addition, two distinct defective swimming behaviors were observed during the swim tunnel experiments in mutant fish (Fig. 5B and Movie S1). First, *col6a1*^*Δex14*^ fish frequently reached the upper part of the tunnel, rather than maintaining position along the Y-axis of the water flow as seen for WT fish. This behavior was interpreted as an increased need to access oxygen at the surface, consistent with previous observations of hypoxic behavior in the mutant line (12) or as an attempt to find a position along the tunnel wall where the flow is slightly lower. However, oxygen consumption was measured over the course of the swim tunnel experiments and showed no significant difference between WT and mutant fish (Fig. S2). The second defect was detected along the X-axis and was interpreted as a sign of fatigue: mutant fish displayed difficulty resisting the water current, moving backward temporarily before returning to the central position. Image analysis of 1-minute swim trajectories at 5 BL·s^-1^, performed as shown in Movie S2, confirmed these observations. Fish tracking, represented as relative displacement within the tunnel, revealed that *col6a1*^*Δex14*^ fish trajectories were more dispersed along both axes as compared to WT fish (Fig. 5C), giving evidence of muscle fatigue and hypoxia-associated behaviors. Quantitative analysis of these defective swimming events, defined as the proportion of frames showing an abnormal displacement along either axis relative to the total number of frames, demonstrated a significantly higher occurrence in mutant as compared to WT fish (Fig. 5D).

## Discussion

The main finding of our study is that the *col6a1*^*Δex14*^ mutation responsible for BM in human induces a reduced expression of functional CaV1.1 and a negative shift of its voltage dependence contributing to an elevated SR Ca^2+^ leak at negative potentials. Throughout this study, we explored all the steps of EC coupling, from excitation to SR Ca^2+^ release. First, we showed that, in absence of ColVI in one-year old BM fish as evidenced by immunofluorescence and western-blot analysis, the amplitude and kinetics of action potentials were not altered by the mutation, indicating that properties of ion channels involved in the firing of action potential were not modified. In contrast, density and voltage sensitivity of the second step of EC coupling, corresponding to the activation of CaV1.1 and resulting in charge movements, were found to be reduced and shifted by 9 mV towards negative membrane potentials respectively. CaV1.1 proteins are mainly anchored in the t-tubule membranes and it might be hypothesized that the observed modifications could be due to a disorganization of this membrane system. However, confocal imaging of fibers stained with di-8anepps and antibodies raised against CaV1.1 showed that density and orientation of t-tubules were not altered in *col6a1*^*Δex14*^ fish. It is thus likely that ColVI deficiency specifically reduces expression of functional CaV1.1 and disrupts its voltage sensitivity. Since CaV1.1 controls in a voltage-dependent manner the opening of the SR Ca^2+^ release channel, it was predictable that the maximal capacity and the voltage-dependence of SR Ca^2+^ release would also be reduced and shifted toward negative voltages respectively. We however did not find any significant change in the maximal amplitude of voltage-activated Ca^2+^ transients and measured a shift of only 5 mV of the voltage-dependence, still significant, but about half of the shift of charge movements, without any change in the inactivation process. In a recent paper, we reported a reduced intramembrane charge movements density in zebrafish muscle as compared to mammalian muscle but likely compensated by a higher efficacy of coupling between CaV1.1 and SR Ca^2+^ release channel since the maximal amplitude of voltage-activated Ca^2+^ transients was found to be comparable to the one recorded in mammalian muscle (14). It thus can be suggested that the changes in charge density and voltage-dependence of CaV1.1 could be dampened following coupling with SR Ca^2+^ release because of this elevated coupling efficiency, explaining in this way the unchanged SR Ca^2+^ release capacity and the lesser voltage shift of SR Ca^2+^ release activation.

Under physiological conditions of activation of the muscle fiber, t-tubular depolarization induced by action potentials always maximally activates SR Ca^2+^ release. Since the maximal capacity of SR Ca^2+^ release was not changed in *col6a1*^*Δex14*^ fish muscle, we envisioned that the hyperpolarizing shift of activation of CaV1.1 and of SR Ca^2+^ release should not greatly impact muscle function during activation. However, CaV1.1 is known to exert an active control on the SR Ca^2+^ release channel over the whole voltage range from resting membrane potential values to values inducing maximal Ca^2+^ release (15). This led us to explore SR Ca^2+^ release capacity at voltages closer to resting membrane potentials that do not elicit detectable intracellular Ca^2+^ changes in response to short duration pulses. We showed that, in response to 10-s duration voltage pulses, the magnitude of Ca^2+^ release was larger for depolarizations from -80 to -60 mV in *col6a1*^*Δex14*^ fibers as compared to WT fibers. These data demonstrate that the shift of voltage-dependence of CaV1.1 activation occurs in mutant fibers over the whole voltage range including membrane potentials close to resting values and may give rise to an elevated SR Ca^2+^ efflux at resting membrane potentials. Additionally, we observed that the basal Ca^2+^ remained at higher levels between voltage pulses in *col6a1*^*Δex14*^ fibers, possibly caused by an increased SR Ca^2+^ efflux at -90 mV. Moreover, when we looked at Ca^2+^ sparks, we measured a significant higher relative area displaying Ca^2+^ sparks in *col6a1*^*Δex14*^ fibers, likely reflecting an elevated SR Ca^2+^ efflux in resting fibers. However, RyR_1_is not the only SR Ca^2+^ release channel present in zebrafish muscle. Another isoform, RyR_3_, has indeed been shown to be localized in the parajunctional SR (18). Silencing RyR_3_by morpholino injection in zebrafish larvae was found to cause a dramatic reduction in Ca^2+^ sparks, suggesting that RyR3 is the main contributor to the SR Ca^2+^ flux underlying Ca^2+^ sparks (18). RyR_3_ opening is considered to be not under the direct control of CaV1.1 but activated by Ca^2+^ flowing through RyR1 *via* a Ca^2+^-induced Ca^2+^ release process as proposed for frog muscle (19). In these conditions, the higher frequency of Ca^2+^ sparks that we observed cannot be directly the consequence of a shift of voltage-dependence of CaV1.1, but, indirectly, could result from a higher efflux of Ca^2+^ through RyR_1_ due to this shift, which in turn over-activates RyR_3_ channels *via* a Ca^2+^-induced Ca^2+^ activation process. Hence, our data lead us to propose that the BM mutation promotes an exacerbated resting SR Ca^2+^ efflux by disrupting the voltage-dependent CaV1.1-gated RyR_1_ activity. Likewise, an evidence supporting this assumption is that intracellular Ca^2+^ increase provoked by inhibition of the mitochondrial F1-F0 ATPase was found to be larger and occurred earlier in muscle from the COLVI-deficient *Col6a1*^*-/-*^ as compared to WT mice. But above all, this Ca^2+^ increase was suppressed by dantrolene, a blocker of the SR Ca^2+^ release channel (10), suggesting the involvement of an elevated SR Ca^2+^ efflux in this Ca^2+^ increase. An elevated resting SR Ca^2+^ efflux has been described in a number of muscle disorders (20, 21). As early observed in Duchenne muscular dystrophy, this elevated SR Ca^2+^ leak is thought to increase protein degradation, mitochondrial dysfunction, and to provoke SR dilation (22), a change in SR morphology that is also observed in BM fish at early stage (12), leading to progressive muscle fiber weakness and degeneration (23). Accordingly, our force measurement experiments showed reduced trunk muscle strength in mutant fish leading to reduced maximal swimming velocity, increased muscle fatigue and hypoxia-associated behaviors in swim tunnel experiments. These data demonstrate that BM mutation does induce muscle weakness in fish as observed in the human disease, likely caused by a pathogenic elevated SR Ca^2+^ efflux.

Although the t-tubule structure is not altered in mutant fish, our CaV1.1 immunolabeling data indicate that the mean distance between fluorescent foci, corresponding to locations of CaV1.1s clusters facing RyR_1_s at the level of triads, is very significantly shorter in mutant fish. This points out mis-localization of CaV1.1s resulting from ColVI deficiency in mutant fish that could explain the observed dysfunction of CaV1.1. Additionally, CaV1.1 proteins are anchored in the membrane of t-tubules that penetrate radially the muscle fiber surrounding myofibrils, so that fluorescent CaV1.1 foci detected with confocal microscopy correspond to CaV1.1 clusters distributed in the same transversal plane separated by distance coinciding with the 1-2 µm range of diameter of myofibrils. Consequently, it could be also envisioned that the shorter distance between foci results from reduced diameters of myofibrils in mutant fish. Such a reduced diameter of myofibrils may be issued from a muscle degenerative process that could further explain the observed reduced force output. Along the same line, adult *col6a1* null zebrafish were found to display defective muscle organization and impaired swimming capabilities (7).

ColVI has been shown to interact with a number of molecules in the extracellular matrix, but the molecular link allowing ColVI deficiency to be transduced into muscle fiber dysfunction remained so far elusive (5). Our study gives evidence that this link, which should necessarily be trans-membranous, is CaV1.1. In contrast to cardiac cell t-tubules, in which ColVI has been found to be located (24), we did not observe ColVI immunoreactivity within the skeletal muscle t-tubules neither of whole mount zebrafish larvae nor of frozen sections of juvenile and adult fish. The diameter of skeletal muscle t-tubule is theoretically large enough for one ColVI filament (6 to 10 nm) to fit into the t-tubule lumen (20 to 40 nm). However, ColVI is generally associated to the muscle basement membrane in skeletal muscles which is about 70 nm thick, 140 nm if we consider that the sarcolemma is invaginated with its basement membrane to form the t-tubules. Additionally, ColVI being produced by fibroblasts in skeletal muscle, it is very unlikely that the ColVI tetramers diffuse to the t-tubules to form filaments that is the last step of ColVI biosynthesis. A direct interaction between ColVI and CaV1.1 is thus very unlikely. But, as proposed by Zanotti et al. (25), we suggest that ColVI deficiency in BM leads to instability in the basal lamina, here evidenced by a change, not only in the ColVI immunoreactivity in BM fish but also in the basement membrane marker laminin, and weakening of the extracellular matrix mechanical support to muscle membrane. Basement membrane plays an important role in providing physical and biochemical cues to the overlying cells (26). It can thus be hypothesized that instability of basal lamina, possibly amplified by cycles of contraction and relaxation and specifically at the level of the t-tubules where it overhangs openings (27), disrupts expression and function of CaV1.1s anchored in the t-tubule membrane. The mechanical properties of ColVI filaments also help stabilize the cell membrane (5). Consequently, ColVI deficiency can alter both basement membrane biomechanics and sarcolemma stability, which in turn compromises the integrity of transmembrane proteins, specifically CaV1.1, which is expressed at extremely high density in t-tubules.

In conclusion, by using a unique zebrafish model of BM, our work led to unravel CaV1.1 as the molecular link transducing ColVI deficiency into alteration of intracellular Ca^2+^ handling and the resulting muscle degenerative process. This study should open new avenues for the treatment of BM using therapeutics targeting CaV1.1.

## Materials and Methods

### Zebrafish maintenance and ethics statement

Wild-type zebrafish (AB/TU) and *col6a1*^*Δex14-/-*^ line (BM fish model created and characterized at larval stage in^12^) maintenance and embryo collection were performed at the zebrafish PRECI facility (Plateau de Recherche Expérimentale de Criblage In vivo; UMS CNRS 3444 Lyon Biosciences, Gerland) in compliance with French government guidelines (see Study approval). Embryos obtained from natural spawning were raised following standard conditions. Developmental stages are expressed in hours, days or weeks post-fertilization (hpf, dpf or wpf, respectively). Equal amount of female and males were used in the study. When larvae were used at two weeks post-fertilization (muscle force analysis), sex determination could not be performed, as at this stage fish lack distinguishable sexual characteristics.

### Generation of guinea pig anti-ColVI antibodies

The recombinant zebrafish α1(VI) C-terminal polypeptide was obtained and purified as described by Tonelotto et al. (28). Purified recombinant α1(VI) C-terminal polypeptide was used for guinea pig immunization, and the obtained antiserum was purified by antigen-affinity chromatography (Covalab, Lyon). The specificity of purified antibodies was determined by ELISA binding assay and immunoblotting and western blot.

### Immunostaining

6-month and 1-year old fishes were terminally anesthetized using 0.2% MS-222 (tricaine methanesulfonate, Sigma-Aldrich). For immunostaining of frozen sections, the trunk was dissected with a razor blade and were fixed overnight at 4°C in 4% PFA and incubated in 30% sucrose in PBS. They were then embedded in OCT freezing medium, and snap-frozen in cold isopentane; 30 µm cryosections were obtained using a Leica cryostat and stored at -80°C until use. For immunostaining, frozen sections were rehydrated for 5 min in PBS and incubated 30 min in blocking buffer (1% BSA, 3% goat serum in PBS). Primary antibodies were then incubated overnight at 4°C. After washes in PBS, secondary antibodies, and Rhodamine phalloidin when indicated, were incubated for 1 h at room temperature. After a rapid wash in PBS, nuclei were stained for 5 min with 1.5 µg/mL Hoechst 33258 (Sigma-Aldrich). Samples were washed and mounted in DAKO mounting medium and stored at 4°C until imaging. Primary and secondary antibodies and Rhodamin phalloidin were used at stated dilutions in blocking buffer: guinea-pig anti-ColVIα1 antibodies (1:2000); rabbit anti-laminin antibody (1:500; Sigma-Aldrich); mouse anti-CaV1.1 (mAb427, Chemicon International); goat anti-guinea pig, anti-rabbit or anti-mouse IgG coupled to AlexaFluor-488 or AlexaFluor-546 (1:500; Invitrogen); and phalloidin-647 (1:80; Sigma-Aldrich). For immunostaining of isolated muscle fibers, trunk muscle of the dorso-caudal region of 1-year old fish were dissected in Tyrode solution and then fixed overnight at 4°C in 4% PFA and processed to immunostaining as described above. Samples were observed with a Zeiss LSM 780 spectral confocal microscope (Zeiss, Oberkochen, Germany).

### Protein extraction and western blot analysis

One-year old fish were euthanized as indicated above. Yolks and head were discarded, the skin was peeled and skeletal muscles were carefully dissected away from the bones in bulk. Samples were snap frozen and stored at -8O°C until further use. Total protein extract was prepared by incubating muscle tissue in extraction buffer containing 50 mM Tris–HCl (pH 6.8), 5% glycerol, 1% SDS, 4 M urea, 50 mM DTT, and 0.1% bromophenol blue at a ratio of 1:20 (mg tissue/μL buffer). Samples were kept on ice and homogenized using a mortar and pestle, followed by repeated syringe aspirations, centrifuged to remove pellet, and the resulting supernatant was heated at 95 °C for 3 min. For Western blot analysis, aliquots of total protein were separated by SDS–PAGE and transferred onto PVDF membranes. Membranes were blocked and incubated with guinea pig anti– ColVI antibodies and mouse monoclonal anti-Vinculin (1:8,000; V4505, Sigma-Aldrich) as a loading control. Immunoreactive bands were detected using HRP-conjugated secondary antibodies and the ImmunStar WesternC substrate (Bio-Rad). The band intensities were quantified by densitometry using ImageJ software, and protein levels were normalized to loading control (Vinculin).

### Isolation and preparation of zebrafish skeletal muscle fibers

Male 10-13-month-old zebrafish were euthanized with 0.3g/L tricaine (MS222, Sigma-Aldrich), decapitated, and skin was removed. Dorsal trunk muscle of around 3 mm width located just under the dorsal fin was removed and incubated for 40 minutes at 37°C in the presence of Tyrode solution containing 2 mg/mL of collagenase (Sigma-Aldrich, type 1). Single intact muscle fibers were then released by gentle mechanical trituration of the enzyme-treated muscles in a glass-bottomed experimental chamber, in the presence of a culture medium containing 2 % Matrigel (Sigma). A previous study indicated that muscle fibers isolated with this procedure were fast type (14).

### T-tubules membrane labelling

For imaging the t-tubule network, isolated zebrafish muscle fibers were incubated for 1h in the presence of 10 μm di-8-anepps in Tyrode solution. Di-8-anepps fluorescence was measured with 488 nm excitation and collected above 505 nm using a confocal microscope equipped with a x63 oil immersion objective and a Zeiss LSM Exciter. To quantify t-tubule organization and sarcomere length, di-8-anepps fluorescence was imaged from 2 distinct longitudinal locations in the fiber, each comprising a minimum of 60 µm-length and the total width. T-tubule organization resembles that in mammalian cardiomyocytes, allowing image analysis with the Image J TTorg plugin based on fast Fourier transformation of images and designed for a reliable and unbiased quantitative analysis of t-tubule organization (29). T-tubules regularity was quantified by the T-tubule power parameter and the mean longitudinal spacing of t-tubules corresponded to the calculated period of di-8-anepps fluorescence.

### Electrophysiology

Prior to trituration, the bottom of the experimental chamber was covered with a thin layer of silicone grease. This enabled single fibers from zebrafish to be covered with additional silicone so that a 50-100 μm-long portion of the fiber extremity was left out, as previously described (30). The culture medium solution was replaced by the extracellular solutions (see Solutions). The tip of a glass micropipette filled with an intracellular-like solution containing a Ca^2+^-sensitive dye (see intracellular Ca^2+^ measurements) was inserted into the silicone-embedded fiber portion. The silver-chloride wire inside the micropipette was connected to an RK-400 patch-clamp amplifier (Bio-Logic, Claix, France) used in whole-cell voltage-clamp or current-clamp configuration. Command voltage or current pulse generation was achieved with an analog-digital converter (Digidata 1322A, Axon Instruments, Foster City, CA) controlled by pClamp 9 software (Axon Instruments). The tip of the micropipette was gently crushed against the bottom of the chamber to reduce the series resistance and to allow internal dialysis of the fiber. Current and voltage changes were acquired at a sampling frequency of 10 and 50 kHz respectively. All experiments were performed at room temperature (20–22°C).

Charge movement currents were calculated using conventional procedures consisting of subtracting scaled control current recorded in response to a 10-mV hyperpolarizing pulse from the current elicited by test depolarizing pulses of identical duration and of various amplitudes from a holding potential of -80 mV. Charge movement was quantified by integrating the transient outward current after the onset of the test pulse (Q_on_) and subsequently normalized to cell capacitance (nC/μF).

### Intracellular Ca^2+^ measurements

#### - Intracellular Ca^2+^ measurements in response to voltage changes

Prior to voltage clamp, the indo-1 dye, diluted at a concentration of 0.2 mM in an intracellular-like solution (see solutions), was dialyzed into the fiber cytoplasm through the microelectrode inserted through the silicone, within the insulated part of the fiber. Intracellular equilibration of the solution was allowed for a period of 20 min before initiating measurements. Indo-1 fluorescence was measured on an inverted Nikon Diaphot epifluorescence microscope equipped with a commercial optical system, allowing the simultaneous detection of fluorescence at 405 nm (F405) and 485 nm (F485) by two photomultipliers (IonOptix, Milton, MA, USA) upon 360 nm excitation. Background fluorescence at both emission wavelengths was measured next to each fiber tested and was then subtracted from all measurements. Fluorescence signals were acquired at a sampling frequency of 50 kHz. The standard ratio method was used with the parameters: R=F405/F485, R_min_, R_max_, Kd and β having their usual definitions. Results were expressed in terms of indo-1 % saturation. In cells values for R_min_, R_max_ and β used were 0.3, 1.61 and 2, respectively. Kd was assumed to be 350 nM. No correction was made for indo-1–Ca^2+^ binding and dissociation kinetics. The displayed Ca^2+^ signal traces were smoothed using adjacent averaging of a number of datapoints specified in figure legends.

#### - Intracellular Ca^2+^ measurements around resting potentials

The non-ratiometric Fluo-4 dye, diluted at a concentration of 0.1 mM in an intracellular-like solution (See solutions), was dialyzed into the fiber cytoplasm through the microelectrode during 20 min before initiating recordings. Fluo-4 fluorescence was measured with 488 nm excitation and collected above 505 nm on a confocal imaging microscope conducted with a Zeiss LSM Exciter and equipped with a x63 oil immersion objective. Cytosolic Ca^2+^ was recording by measuring the changes of Fluo-4 fluorescence elicited by 25 consecutive confocal frames every 500 ms during 10-s voltage clamp depolarizing pulses of increasing amplitude from a holding potential of -90 mV. Fluo-4 Ca^2+^ transients were expressed as F/F_0_, where F_0_ is the basal fluorescence measured at - 90 mV.

#### - Measurement of Ca^2+^ sparks

Isolated muscle fibers were incubated for 30 min in the presence of 10 μmol/L Fluo-4 acetoxymethyl ester (AM). After 4 times washing with Tyrode solution, 40 consecutive confocal frames of fluo-4 fluorescence were acquired in each fiber at 1 Hz frequency. To quantify the Ca^2+^ sparks activity, images in the stack were smoothed and the standard deviation of fluorescence intensity at each pixel position, along the stack, was calculated. The 10% largest values in the standard deviation image were removed to calculate the mean standard deviation of silent areas. The active area was then defined as pixel positions exhibiting at least 1.5 × larger values of standard deviation than the mean standard deviation value from silent areas. This active area was expressed as percentage of total analyzed area.

#### Measurement of trunk muscle force

Procedures and measurements were performed as previously reported (31). Two wpf fish were briefly placed in a beaker containing 0.168 mg/mL of tricaine (3-amino benzoic acidethylester, Sigma-Aldrich, A5040). The head was crushed and fish were transferred to a Tyrode solution. Under binocular control, fish head and tail extremities were glued with surgical glue (Histoacryl, B Braun) on homemade thin aluminum foils pierced with a small hole allowing to attach the head portion of the fish to the arm of an AE801 force transducer and the tail portion to a fixed pin. Fish were stimulated by field electrodes placed on either side of the fish using a Harvard apparatus 6002 stimulator. In each fish, the voltage of 0.5 ms duration pulses was increased to give maximal twitch response and the voltage used for measuring maximal force was set at 20% above this maximal value. The fish was then gradually stretched on the side of the transducer moved with a micromanipulator until maximal twitch force was obtained. The force signal was recorded at 10 kHz sampling frequency using the WinWCP software (Strathclyde University, UK) driving an AD converter (National Instruments, USA).

#### Measurement and image analysis of swimming performance

Swimming performance was measured using a swim tunnel (Loligo Systems, Denmark). Male and female fish at one year of age were used indiscriminately for analysis and anesthetized with tricaine for biometry measurements (Table S1). Only females with an abdomen full of eggs, which could impair swimming capacity, were excluded. The tunnel is composed of a swimming chamber, where the fish to be tested is placed, a motor fitted with three-bladed propeller to control water flow and a honeycomb placed at each side of the swimming chamber to laminarize the water flow. Before being placed into the swim tunnel, fish were starved 24 hours. Then, the fish was submitted to a short acclimation swimming period (0.5 body length (BL).s^-1^) prior to starting the swimming challenge. The challenge consists to an increase of the water flow by stage of 1.5 BL. s^-1^ every 15 min. Between each stage, water was refreshed during 3 min. The experiment stopped when the fish did not manage to swim against the current. Then, water flow was reduced to initial value to permit fish to recover. Fish were then removed from the swim tunnel and anaesthetized using tricaine to determine the total length, mass and sex of each individual. Swimming performance was calculated as critical swimming speed (U_crit_) according to the formula: U_crit_ = Ut + t_1_.t^-1^ .U_1_, where Ut (in BL.s^−1^) is the highest velocity maintained for an entire step, t_1_ (in min) is the time spent until the exhaustion of fish at the last step, t (in min) is the swimming period for each step (*i*.*e*. 15 min in the present study) and U_1_ is the increment velocity (1.5 BL.s^−1^) (32).

Fish were detected and their movements were analyzed using MicrobeJ 5.13p (33) a plugin for ImageJ (34). Briefly, all test videos were acquired using the same imaging equipment at 30 frames per second. For fast processing speed, original videos were converted to 8-bit monochrome stack of image slices using ffmpeg (https://ffmpeg.org/) and resized to the resolution of 624×264 pixels for further analysis. Rigid body registration using the StackReg plugin (35) was performed on each stack of images to correct translation and/or angular drift that could have occurred during the acquisition. Fishes were detected by template matching using a 12×12 pixels image of the fish eye, as a kernel. If needed, the fish detections were manually corrected using the editing interface of MicrobeJ. Each fish was tracked using a nearest neighbor algorithm, and the resulting trajectories were analyzed to identify specific behaviors. The occurrence and duration of each specific behavior were recorded for each fish and visualized using an event plot.

### Solutions

For electrophysiology and Ca^2+^ measurements on isolated muscle cells, the dialyzed intracellular solution contained (in mM): 140 K-glutamate, 5 Na_2_-ATP, 5 Na_2_phosphocreatine, 5.5 MgCl_2_, 5 glucose and 5 Hepes adjusted to pH 7.2 with KOH, except for the experiments devoted to the measurement of intramembrane charge movements, for which the dialyzed intracellular solution contained (in mM): 140 Cs-aspartate, 5 Mg-ATP, 1 MgCl_2_, 10 EGTA-CsOH and 5 Hepes, adjusted to pH 7.2 with CsOH. The Tyrode solution used in current-clamp conditions contained (in mm): 140 NaCl, 5 KCl, 2.5 CaCl_2_, 2 MgCl_2_ and 10 Hepes, adjusted to pH 7.2 with NaOH. The extracellular solution used for measurements of charge movements and Ca^2+^ signals in voltage-clamp conditions contained (in mm): 140 TEAmethanesulphonate, 2.5 CaCl_2_, 2 MgCl_2_, 1 4AP, 0.002 TTX and 10 Hepes, adjusted to pH 7.2 with TEAOH. N-benzyl-p-toluene sulphonamide was added in extracellular solutions at 50 μM.

### Statistics

Sample size was estimated according to a statistical analysis consisting on performing a comparison of the means of two unpaired group with a 5 % risk level alpha and a power of 80 % using bilateral test. Experiments were performed using WT and mutant fish drawn from at least three different tanks placed at different positions within the tank holder to minimize location-related variations. The order of treatments and measurements was randomized across experimental days to avoid systematic bias. Randomization was also ensured because fish were obtained from multiple mating pairs of different lineages.

For electrophysiological experiments and intracellular Ca^2+^ measurements performed on isolated muscle fibers (Fig. 2E, 3 and 4), statistical analysis was performed using Microcal Origin and GraphPad Prism. Least-squares fits were performed using a Marquardt–Levenberg algorithm routine included in Microcal Origin. Normality of data distribution was assessed using Shapiro-Wilk test and statistical differences were determined using a nested t-test taking into account the number of fibers from each animal. For data not normally distributed, statistical differences were determined using unpaired Mann Whitney Wilcoxon test. Data are given as means ± S.E.M. Numbers of individual measurements and individual animals used are mentioned in the figure legends. Differences were considered significant when *P* < 0.05. Labels *, **, *** and **** indicate *P* < 0.05, *P* < 0.01, *P* < 0.001 and *P* < 0.0001 respectively.

For figures 1, 2B and C, 5, 6 and S2, the statistical test used, the number of animals used and P values are indicated in the figure legends.

### Study approval

All animal manipulations were performed in agreement with EU Directive 2010/63/EU. The creation and characterization of the *col6a1*^*Δex14-/-*^ line (Bethlem myopathy fish model) was described in (12). The study has been approved by the French Ministère de l’Enseignement Supérieur de la Recherche et de l’Innovation with the number APAFIS#28791-2020102114261517 v4.

## Acknowledgments

This work was supported by CNRS, Inserm, INRAE, Ecole Normale Supérieure (ENS) de Lyon, Université Lyon 1, Association Française contre les Myopathies (AFM), Agence Nationale de la Recherche (ANR) (FishandCol6 project). RI is the recipient of PhD fellowship funded by the Fondation pour la Recherche Médicale (FRM). SS is the recipient of PhD fellowship funded by the European Union within the Horizon Europe MSCA programme under grant agreement N° 101072766 (CHANGE project). We thank Laurent Gilquin (IGFL, ENS de Lyon) for his help in CaV1.1 distribution quantification.

## Supplemental materials

### Supplementary figures

**Figure S1. Variability in ColVI residual staining in 1-year-old *col6a1***^***Δex14***^ **fish**. Representative confocal images of 1-year old zebrafish muscle cross-sections from 3 independent experiments (1-3) stained with anti-ColVI antibodies. Number of fish per experiment shown on the right.

**Figure S2. Oxygen consumption of *col6a1***^***Δex14***^ **fish during effort**. Measurements of oxygen consumption (MO_2_) during the step protocol of WT and *col6a1*^*Δex14*^ fish. The numbers indicate the number of fish analyzed. WT (n°6-13, Table S1) and *col6a1*^*Δex14*^ (n°1-14, Table S1) were used for the analysis of O_2_ consumption. Data are means ± SEM. Statistical analysis was performed using a two tailed Student’s t-test for paired samples.

### Supplementary table

**Table S1. Biometry of fish used for swimming tunnel step protocol**. M, male; F, female; SL, standard length; TL, total length. The fish WT10 and *col6a1*^*Δex14*^ 4 (lines colored in grey) were excluded from the data analysis due to technical problems during the swim tunnel assays.

### Supplementary movies

**Movie S1**. Swimming tunnel montage of representative 1-min movies of WT (upper row) and *col6a1*^*Δex14*^ fish (lower row) at 5 BL·s^−1^. WT fish number 5, 6, 7, 10 and 13 and *col6a1*^*Δex14*^ fish 1, 2, 7, 8 and 14 (as referred in Table S1) are shown (50 frames/s). Defective swimming behavior along the X and Y axes is indicated by yellow boxes.

**Movie S2**. Interactive swim tracking with corresponding image analysis. The blue line indicates the expected central position of the fish during exercise, while the yellow line shows deviations from the barycenter along the X-axis (upper trace) or Y-axis (lower trace). A representative *col6a1*^*Δex14*^ fish (number 2 in Movie S1, Table S1) is shown as an example.

