## Supplementary material for "A zebrafish model of Bethlem myopathy reveals CaV1.1 as the missing link between collagen type VI deficiency and muscle dysfunction": Figure S1

**Supplemental information**

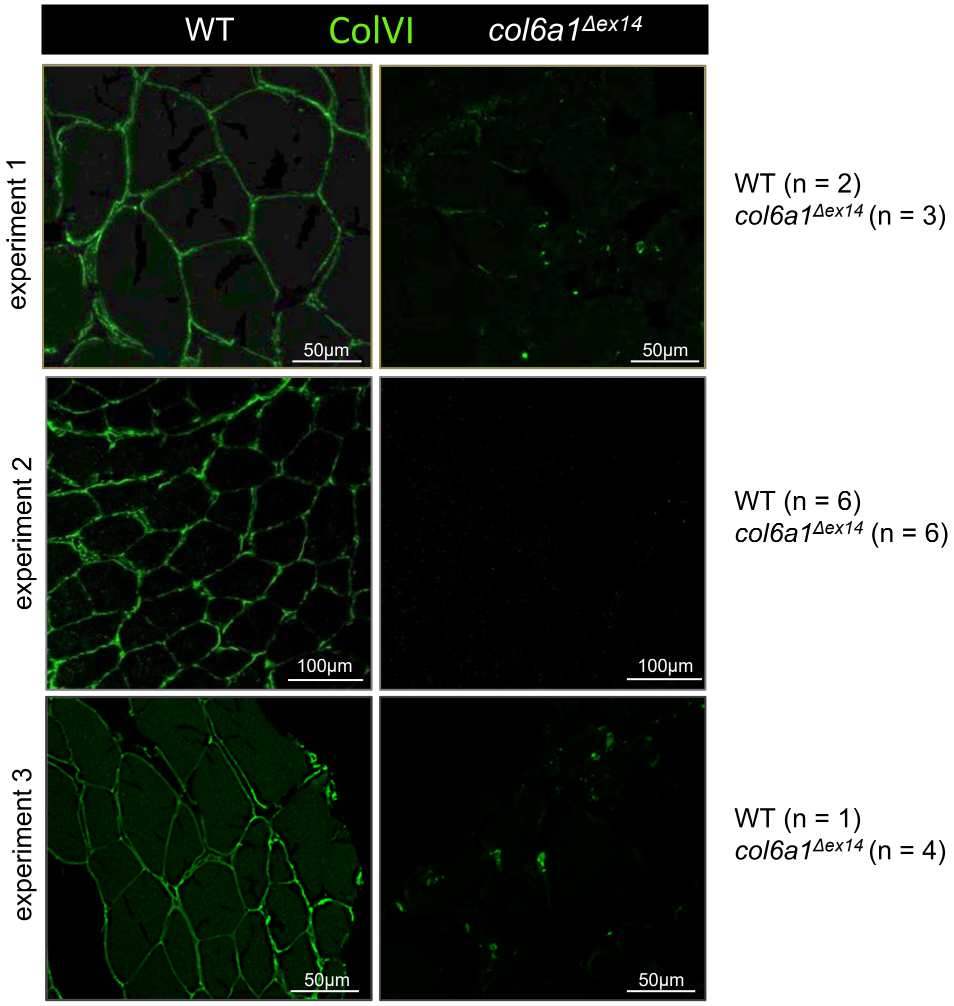


**Figure S1. Variability in ColVI residual staining in 1-year-old *col6a1*$^{\Delta ex14}$ fish.** Representative confocal images of 1-year old zebrafish muscle cross-sections from 3 independent experiments (1-3) stained with anti-ColVI antibodies. Number of fish per experiment shown on the right.
