## Supplementary material for "A zebrafish model of Bethlem myopathy reveals CaV1.1 as the missing link between collagen type VI deficiency and muscle dysfunction": Figure S2

**Supplemental information**

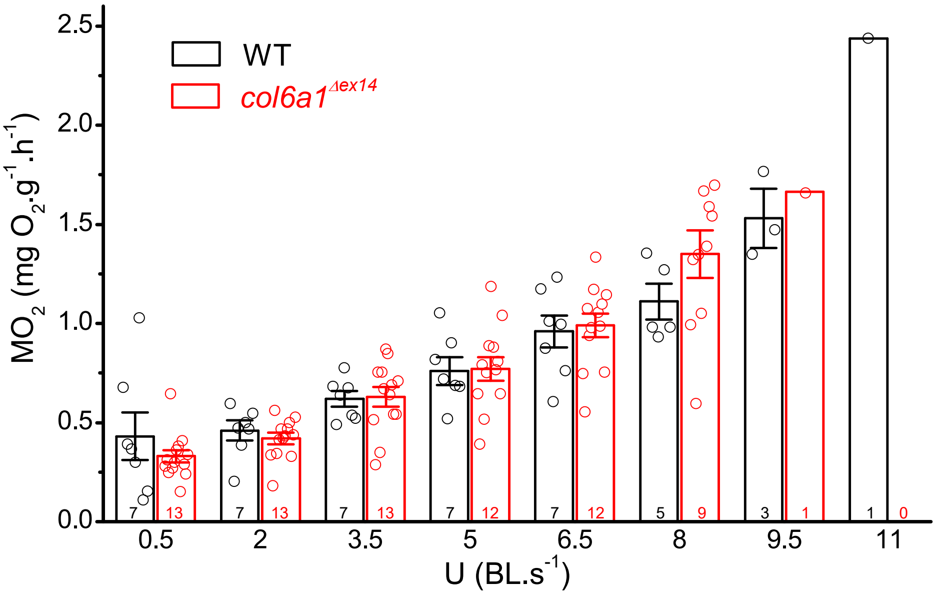


**Figure S2. Oxygen consumption of *col6a1*$^{Δex14}$ fish during effort.** Measurements of oxygen consumption ($MO_2$) during the step protocol of WT and *col6a1*$^{Δex14}$ fish. The numbers indicate the number of fish analyzed. WT (n°6-13, Table S1) and *col6a1*$^{Δex14}$ (n°1-14, Table S1) were used for the analysis of $O_2$ consumption. Data are means ± SEM. Statistical analysis was performed using a two tailed Student's t-test for paired samples.
