## Supplementary material for "A zebrafish model of Bethlem myopathy reveals CaV1.1 as the missing link between collagen type VI deficiency and muscle dysfunction": Table S1

### Supplemental information

| Fish genotype and number | Sex | Weight (g) | Morphometry | | | |
|---|---|---|---|---|---|---|
| | | | SL (cm) | TL (cm) | Thickness (cm) | Height (cm) |
| WT1 | F | 0.631 | 2.9 | 3.4 | 0.2 | 0.9 |
| WT2 | M | 0.501 | 3.1 | 3.7 | 0.3 | 0.7 |
| WT3 | F | 0.354 | 2.6 | 3.2 | 0.3 | 0.8 |
| WT4 | M | 0.664 | 3.3 | 3.9 | 0.3 | 0.8 |
| WT5 | F | 0.730 | 3 | 3.7 | 0.4 | 0.9 |
| WT6 | M | 0.573 | 3.2 | 4 | 0.2 | 0.7 |
| WT7 | F | 0.830 | 3.1 | 3.8 | 0.5 | 1 |
| WT8 | M | 0.430 | 2.9 | 3.6 | 0.3 | 0.7 |
| WT9 | F | 0.593 | 3 | 3.7 | 0.4 | 0.8 |
| WT10 | M | 0.426 | 2.8 | 3.6 | 0.3 | 0.7 |
| WT11 | M | 0.405 | 2.9 | 3.6 | 0.3 | 0.7 |
| WT12 | F | 0.481 | 3.1 | 3.8 | 0.4 | 0.8 |
| WT13 | M | 0.546 | 3.1 | 3.9 | 0.3 | 0.7 |
| *col6a1*$^{\Delta ex14}$1 | M | 0.566 | 3.1 | 3.9 | 0.3 | 0.7 |
| *col6a1*$^{\Delta ex14}$2 | F | 1.030 | 3.2 | 3.9 | 0.4 | 0.8 |
| *col6a1*$^{\Delta ex14}$3 | F | 0.762 | 3.4 | 4.1 | 0.4 | 0.9 |
| *col6a1*$^{\Delta ex14}$4 | M | 0.463 | 3.1 | 3.9 | 0.3 | 0.7 |
| *col6a1*$^{\Delta ex14}$5 | M | 0.846 | 3.3 | 4.03 | 0.4 | 0.7 |
| *col6a1*$^{\Delta ex14}$6 | F | 0.574 | 3.1 | 3.7 | 0.4 | 0.8 |
| *col6a1*$^{\Delta ex14}$7 | F | 0.960 | 3.8 | 4.6 | 0.5 | 1.1 |
| *col6a1*$^{\Delta ex14}$8 | M | 0.582 | 3.2 | 4 | 0.4 | 0.8 |
| *col6a1*$^{\Delta ex14}$9 | M | 0.460 | 3.1 | 3.7 | 0.3 | 0.7 |
| *col6a1*$^{\Delta ex14}$10 | M | 0.533 | 2.9 | 3.8 | 0.3 | 0.7 |
| *col6a1*$^{\Delta ex14}$11 | M | 0.550 | 3.1 | 3.9 | 0.3 | 0.7 |
| *col6a1*$^{\Delta ex14}$12 | M | 0.606 | 3.1 | 3.8 | 0.3 | 0.7 |
| *col6a1*$^{\Delta ex14}$13 | M | 0.562 | 3.1 | 3.9 | 0.3 | 0.7 |
| *col6a1*$^{\Delta ex14}$14 | M | 0.460 | 3.5 | 3.8 | 0.3 | 0.7 |

**Table S1. Biometry of fish used for swimming tunnel step protocol.** M, male; F, female; SL, standard length; TL, total length. The fish WT10 and *col6a1*$^{\Delta ex14}$ 4 (lines colored in grey) were excluded from the data analysis due to technical problems during the swim tunnel assays.
